# Genome assembly and gene expression in the American black bear provides new insights into the renal response to hibernation

**DOI:** 10.1101/316596

**Authors:** Anuj Srivastava, Vishal Kumar Sarsani, Ian Fiddes, Susan M. Sheehan, Rita L. Seger, Mary E. Barter, Selena Neptune-Bear, Charlotte Lindqvist, Ron Korstanje

**Author notes:** Correspondence: Dr. Ron Korstanje, The Jackson Laboratory, 600 Main Street, Bar Harbor, Maine.

## Abstract

The prevalence of chronic kidney disease (CKD) is rising worldwide and 10-15% of the global population currently suffers from CKD and its complications. Given the increasing prevalence of CKD there is an urgent need to find novel treatment options. The American black bear (*Ursus americanus*) copes with months of lowered kidney function and metabolism during hibernation without the devastating effects on metabolism and other consequences observed in humans. In a biomimetic approach to better understand kidney adaptations and physiology in hibernating black bears, we established a high-quality genome assembly. Subsequent RNA-Seq analysis of kidneys comparing gene expression profiles in black bears entering (late fall) and emerging (early spring) from hibernation identified 169 protein-coding genes that were differentially expressed. Of these, 101 genes were downregulated and 68 genes were upregulated after hibernation. Fold changes ranged from 1.8-fold downregulation (*RTN4RL2*) to 2.4-fold upregulation (*CISH*). Most notable was the upregulation of cytokine suppression genes (*SOCS2, CISH,* and *SERPINC1*) and the lack of increased expression of cytokines and genes involved in inflammation. The identification of these differences in gene expression in the black bear kidney may provide new insights in the prevention and treatment of CKD.

## 1. Introduction

The prevalence of chronic kidney disease (CKD) is rising worldwide and currently 10-15% of the global population suffer from CKD and its devastating complications ^1^. Although the adult human kidney has some ability to recover from acute kidney injury through cellular proliferation of the damaged intrarenal tissues, regenerating nephrons through *de novo* nephron development is considered unlikely, as the formation of new nephrons in humans is terminated at the embryonic stages. However, renal regeneration through nephron neogenesis in the event of renal injury has been described in some fish species and the possibility of this happening in mammalian species has not been excluded. Members of the bear family (Ursidae) might be such species and studying how they deal with periods of decreased kidney function during hibernation (biomimicry) could be a new approach to understand kidney disease and develop new treatments^2^.

The American black bear hibernates for up to seven months annually. During this period, they do not eat, drink, urinate or defecate. Bear hibernation is a state similar to prolonged sleep during which body temperature is reduced by 1-8 C ^3^, there is a 20-50% reduction in metabolic rate with a depressed heart rate ^4^, and the volume of urine produced is reduced by 95% ^5^. The small volumes of urine and urea that enter the bladder during hibernation are reabsorbed across the bladder epithelium ^5^, and the urea is recycled for production of new protein ^6^. Throughout hibernation the kidney continues to concentrate urine and produces renin ^7^, erythropoietin ^5^, and vitamin D 1-α-hydroxylase ^8^. Hibernating bears have the ability to prevent azotemia (high levels of nitrogen-containing compounds in the blood, common in human patients with renal function), but the mechanism is unknown.

Understanding these processes could lead to creating novel therapies for treating human conditions related to resistance to the complications of CKD and recovery from acute kidney injury. The studies mentioned above suggest unique kidney features in the American black bear allowing them to endure lower functioning during hibernation and recovery soon after hibernation. They are likely in part encoded in the genome sequence and gene expression patterns unique to the bear. To address this, we have performed high-throughput sequencing of genomic DNA and RNA isolated from kidneys of wild black bears. We generated a *de novo* assembly and annotation of the complete genome and compared transcription profiles of kidneys collected in the spring (within weeks after hibernation) and in the fall (before hibernation).

## 2. Materials and methods

### 2.1. Sample collection, library preparation and sequencing

Bear samples were obtained by hunters during the hunting seasons in Maine. Hunters were asked to participate on a voluntary base and no bears were killed for the specific purpose of this study. All methods were carried out in accordance with relevant guidelines and regulations. DNA was isolated using the DNeasy Blood & Tissue Kit (Qiagen). The whole-genome library was prepared using the KAPA Hyper Prep Kit (Kapa Biosystems, Inc., Wilmington, MA) with a bead-based size selection to select for inserts with an average size of 400 bp and 6 cycles of PCR. Sequencing was done on two 2×125 bp Illumina 2500 lanes. The mate pair library was prepared using the Illumina Nexera Mate Pair Kit (Illumina, San Diego, CA, USA) with a gel-based size selection to select for inserts with an average size of 10 kb and 14 cycles of PCR. Sequencing was done on two 2×100 bp Illumina 2500 lanes. The PacBio library was prepared using the Pacific Biosciences SMRTbell Template Prep Kit 1.0 (Pacific Biosciences, Menlo Park, CA, USA) using the “20-kb Template Preparation Using BluePippin Size-Selection System (15-kb Size cutoff)” protocol obtained from PacBio SampleNet. The BluePippin was set to collect from 7-50 kb. Sequencing was done on 23 PacBio SMRT cells. For RNA-Seq, hunters collected kidney samples and stored them in liquid nitrogen. Samples were first homogenized in Trizol (Invitrogen) and isolated using the miRNeasy Mini kit (Qiagen), according to manufacturers’ protocols, including the optional DNase digest step. Quality was assessed using an Agilent 2100 Bioanalyzer instrument and RNA 6000 Nano LabChip assay (Agilent). We prepared total RNA for sequencing using the Illumina TruSeq methodology (TruSeq Stranded Total RNA LT Sample Prep Kit with Ribo-Zero Gold). The first step involves the removal of ribosomal RNA (rRNA) using biotinylated, target-specific oligos, combined with Ribo – Zero rRNA removal beads. After individual samples were bar-coded they were pooled and sequencing was done on two 2 x 100 bp Illumina 2500 lanes. All raw data has been submitted to the NCBI’s Sequence Read Archive (http://www.ncbi.nlm.nih.gov/sra) under accession number SRP075217.

### 2.2. Sequence assembly

All data was subjected to quality control check ^9^ and samples with base qualities greater ≥ 30 over 70 % of read length were used in the downstream analysis. KmerFreq_HA v2.0 and Corrector_HA v2.01 tools of SOAPec_v2.01 ^10^ were further used to perform K-mer frequency generation and error correction of paired-end and mate-pair data, respectively. SOAPdenovo-127mer (v2.04) ^10^ was used to perform the contig and scaffold assembly from paired end and mate-pair libraries (avg_ins = 350 and 10K for paired-end and mate-pair, respectively). After the assembly, gaps in scaffolds were closed with the GapCloser tool (v1.12) ^10^ (with – l 125) of the soap module. Smaltmap, perfectfrombam and pipeline tools of the reapr module (v 1.0.17) ^11^ was used to recognize the errors in the assembly by re-mapping of paired-end and mate-pair data to the de novo assembled genome. The assembly was broken at potential misassembled points. The broken assembly was further used as an input for the jelly tool of the PBSuite ^12^ (v15.2.20) with blasr (v1.3.1) at parameters [-minMatch 8-minPctIdentity 80-bestn 1-nCandidates 20-maxScore-500]. This tool was used to upgrade the existing Illumina assembly with low-pass PacBio data. Repeat masking was performed by downloading the Repbase TE library from the repbase server (http://www.girinst.org/server/RepBase/). To identify known TE elements, we used repeat masker and repeat protein mask software in the Repeat Masker package (www.repeatmasker.org), which identifies TEs by aligning the genome sequence to a defined TE database. Tandem repeats were predicted using TRF ^13^ by using the default parameters “Match = 2, Mismatch=7, Delta=7, PM=80, PI=10, Minscore=50 and MaxPeriod=12. The completeness of the assembly was estimated by using CEGMA ^14^. The assembly was screened against a collection of 248 universal eukaryotic single-copy genes. Core eukaryotic gene datasets were downloaded and a blast database was made from the assembly before running CEGMA. This Whole Genome Shotgun project has been deposited at DDBJ/ENA/GenBank under the accession LZNR00000000. Version LZNR01000000 is described in this work. It is also available at ftp://ftp.jax.org/maine_blackbear_project/

### 2.3. Annotation

The black bear genome assembly was annotated using Comparative Annotation Toolkit (CAT--https://github.com/ComparativeGenomicsToolkit/Comparative-Annotation-Toolkit). CAT uses whole genome alignments generated by progressiveCactus ^15^ to project gene annotations from a high quality reference genome on to one or more target genomes. This process leverages previously curated annotation sets to rapidly construct a set of orthologs in the target genomes. After transcript projection ^16^, the protein coding projections are provided to the *ab-initio* gene finding tool AUGUSTUS ^17^ with additional extrinsic information derived from RNA-seq. AUGUSTUS then enforces a coding gene model that allows for shifts in splice sites in order to maintain frame. This process can also rescue exons that did not align in the whole genome alignment. CAT also performs true *ab-initio* gene prediction by using a new method of running AUGUSTUS called Comparative Augustus, or AugustusCGP ^18^. This parameterization simultaneously predicts genes in all species in a progressiveCactus alignment, using RNA-seq data in one or more species to help guide the annotation process. After projection, clean-up and *ab-initio* prediction, CAT combines these separate transcript sets into one final gene set through a consensus finding process. For orthologous transcripts, the transcript with the best fidelity to the reference with the best extrinsic support is chosen. In the case where multiple paralogous projections are found, CAT selects the most likely ortholog through a combination of splice junction fidelity, synteny and alignment identity. Finally, *ab-initio* predictions from AugustusCGP are evaluated for providing new information. If significant overlap with a orthologous projection is found, and the transcript provides a new splice junction or exon, then it is included as a new isoform of the ortholog. Otherwise, the locus is considered a candidate for a novel gene, often a gene family expansion. In addition to an annotation set on target genomes, CAT produces a UCSC Assembly Hub ^19^. This assembly hub has tracks for the raw transMap projections, the post-filtering projections, the various modes of AUGUSTUS employed, the final consensus annotation set, as well as the input RNA-seq information including a filtered splice junction track. All annotations are stored in a modified bigBed format with a wide variety of various additional features annotated on each entry, which can be accessed from the details page for that entry. Additional features include RNA-seq support for specific splice junctions and binary classifiers such as having a frame-shifting indel relative to the source transcript. In addition to the genome-specific tracks, an alignment track (snake track) shows the cactus alignment between the current species and other species. Browsing this assembly hub provides the opportunity to visualize the relationship between all aligned species and the various ways the transcript projections were filtered and combined with *ab-initio* predictions. Assembly hub can be visualized by visiting UCSC “Track Data Hubs” (https://genome.ucsc.edu/index.html) and then adding URL ftp://ftp.jax.org/maine_blackbear_project/assemblyHub/hub.txt under the “My Hubs” tab. For the black bear project, the cactus alignment generated contained black bear, horse (equCab2), dog (canFam3), polar bear (GCA_000687225.1), elephant (loxAfr3), human (hg38) and mouse (mm10) using the following guide phylogenetic tree: ((Human:0.145908,Mouse:0.356483)human_mouse_anc:0.020593,(Horse:0.109397,(Dog:0.052458,(Polar_bear:0.01,Maine_black_bear:0.01)bear_anc:0.08)dog_bear_anc:0.069845)horse_dog_bear_anc:0.043625)root; CAT was run using the Dog Ensembl V87 annotation set as the reference. RNA-seq was obtained from SRA for dog (SRR2960309, SRR2960315, SRR2960317, SRR2960319, SRR2960320, SRR2960321, SRR2960328, SRR3727716, SRR3727717, SRR3727718, SRR3727719, SRR3727720, SRR3727722, SRR3727723, SRR3727724, SRR3727725, SRR5018836) and for polar bear (SRR950076, SRR950074) and aligned to their respective genomes. Additionally, RNA-seq was generated for Maine black bear as part of this project.

### 2.4. Gene expression analysis and RNA editing

Samples were aligned to the soft masked Bear Assembly using STAR aligner (v2.5.3) (-- sjdbOverhang 75-- quantMode GeneCounts--twopassMode Basic) with known annotation. STAR provided the expression count matrix was used for differential expression analysis with DESeq2. The genes with FDR < = 0.05 were considered as differentially expressed between spring and fall Samples. STAR generated bam were further processed by picard MarkDuplicates and GATK SplitNCigarReads to remove the duplicates and split reads into exonic segments and hard clip any reads hanging into the intronic region, respectively. The final bam of all twelve samples were used to perform the join variant calling by HaplotypeCaller (v3.4-0). The output of HaplotypeCaller was further processed to identify the sites showed alternate allele in all twelve samples or that were genotype as reference in all six samples of one season and as alternate allele in the other season. We further filtered variants to set of canonical editing sites (and their reverse complementary). RNA editing prediction tool-REDItools was used to confirm the filtered sites, strand identification and its count was used as final count to estimate the editing frequency. These sites were subjected to confirmation in DNA-seq data generated for six samples. The whole genome sequence data of these bears were subjected to quality trimming and aligned by bwa-mem (v0.7.9a). The alignments were converted to bam and processed by Picard-MarkDuplicates to remove the duplicates from the data. Afterwards, the Genome Analysis tool kit module IndelRealigner was used to pre-process the alignments. The realigned bam file was processed to identify the coverage at potential editing sites. The sites with at least one DNA-seq sample support were included in final set for experimental validation. Gene expression profiles from embryonic mouse kidneys were downloaded from NCBI GEO (https://www.ncbi.nlm.nih.gov/geo/query/acc.cgi?acc=GSE3808) ^20^. Six samples representing three time points (two replicates each) were used in the analysis. For the analysis, E12.5, E13.5 and E16.5 were considered early & late age samples, respectively. GEO samples accessions of array data are GSM87387, GSM87388, GSM87389, GSM87390, GSM87391, and GSM87392. Samples were normalized using the affy package ^21^ and GenBank accession for the probs (Affymetrix Mouse Expression 430A Array) were downloaded from GEO (https://www.ncbi.nlm.nih.gov/geo/query/acc.cgi?acc=GPL339). We extracted the MGI Gene IDs corresponding to GenBank accession using mouse genome informatics database (http://www.informatics.jax.org), performed the principal component analysis (PCA) on the mouse array data and created a PCA biplot. To map the differentially expressed (DE) bear genes on the PCA biplot, we first extracted the corresponding dog gene Ensembl id from the bear generic feature format file (gff) and extracted the Ensembl id of mouse gene orthologs of dog from biomart ^22^. Only genes with homology type: “ortholog_one2one”, orthology confidence:high and Query:target & target:query identity (≥50 percent) between dog and mouse were selected (153 genes remaining at the end) for mapping. Finally, we extracted the MGI:ID for the mouse ensembl id and mapped it on the mouse PCA biplot to create an eggogram.

## 3. Results

### 3.1. Sequencing and Assembly of the Black Bear Genome

The ability to explore the black bear genome for unique features and to facilitate gene expression analysis depends on an assembled and well annotated genome. High-throughput sequencing of the black bear genome has been previously reported ^23^, but only using Illumina short-read sequencing to a ∼30x average depth of coverage. We improved this coverage by extracted DNA from a single male Maine black bear and sequencing it using three methods: paired-end sequencing and mate-pair sequencing using the Illumina HiSeq 2500 platform and single molecule sequencing using the PacBio RSII system (Table S2 and S3). Raw reads were assembled into 113,759 scaffolds and contigs ≥1000 bases with an N50 length of 190 Kb, totaling 2.59 Gb in length (Table S4). This is slightly larger than the estimated size of the panda bear assembly (2.46 Gb) ^24^ and the polar bear assembly (2.53 Gb) ^23^.

We estimated the quality of the assembled sequence by mapping all paired-end reads back to the assembled genome with Burrows-Wheeler Aligner^25^ to determine the mappability and median coverage of the assembled genome. Approximately 90% of reads mapped back to the assembly with a mapping quality of ≥ 30 and with a median coverage of 43x. The peak sequencing depth was 50x, and more than 20 reads covered over 85% of the assembled sequences (Fig. 1a). The completeness of our assembly was estimated using CEGMA by screening against 248 highly conserved core eukaryotic genes. Our black bear assembly covers 212 out of 248 genes completely and 241 out of 248 genes partially. Figure 1b shows this 85.5% completeness compared to 88.7% for the panda bear assembly and 90.7% for the polar bear assembly, which are all comparable. GC content in mammals is correlated with a number of genomic features that are functionally relevant, for example, gene density, transposable element distribution, and methylation rate. The GC-content distribution in the black bear covers a narrower range compared to the panda bear and the polar bear (Fig. 1c). This might be a consequence of the lower amount of repetitive sequences that is present in the black bear genome (Fig. 2). Thus, we were able to establish a genome assembly of good coverage and quality for the black bear, comparable to polar bear and panda bear.

**Fig. 1.**
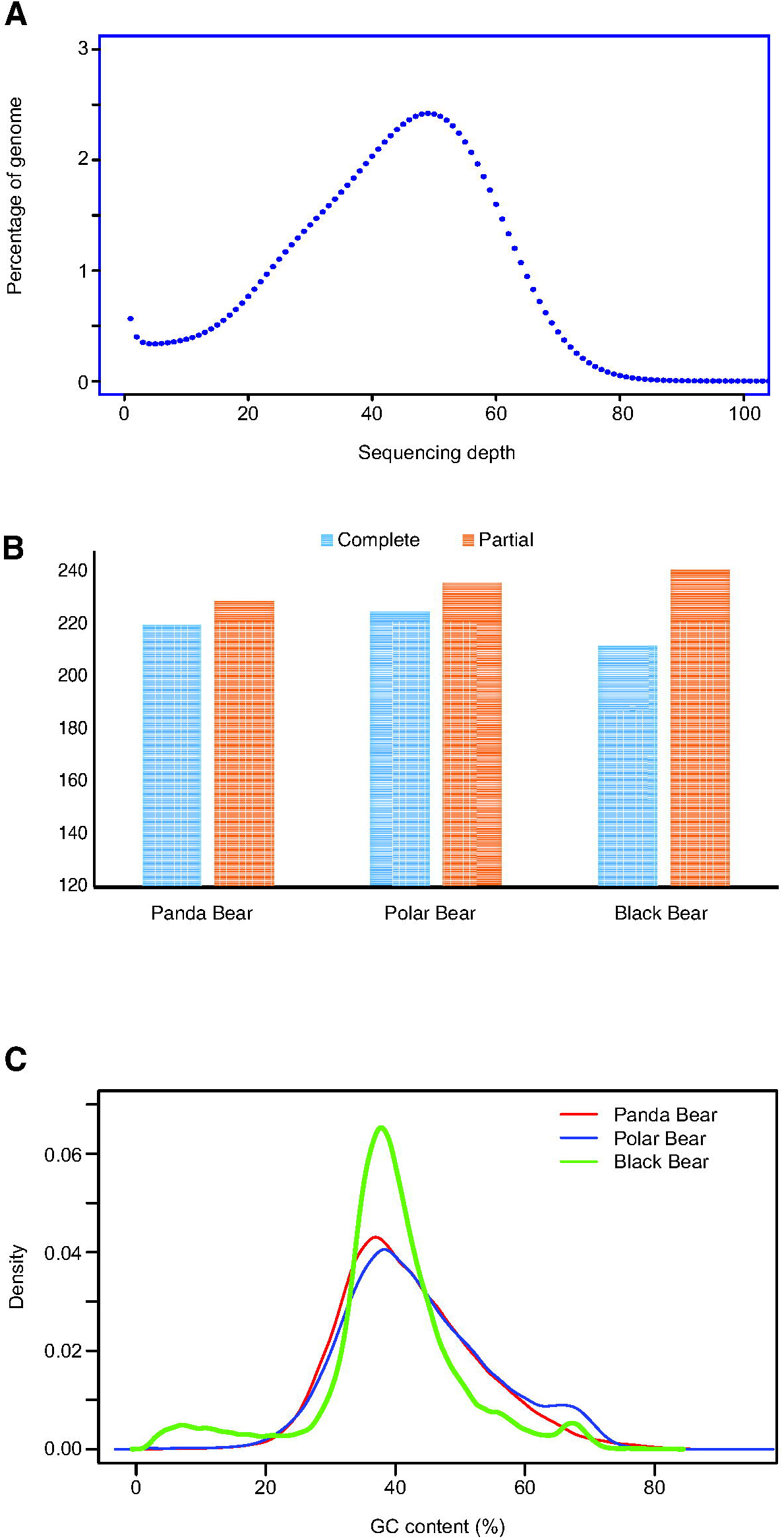
Sequencing depth and comparison of several characteristics of the black bear, panda, and polar bear genome assemblies. (A) Distribution of sequencing depth of the assembled genome (B) Completeness and contiguity of the assembly was estimated using CEGMA by screening against 248 universal eukaryotic single-copy genes. Results for the black bear (85.5% completeness) were comparable to the panda (88.7%) and polar bear (90.7%). (C) Density plot of the %GC content in the bear genomes comparing the black bear (solid line) with the panda (dashed line) and polar bear (dotted line). The narrower distribution in the black bear might indicate fewer repetitive sequences.

**Fig. 2.**
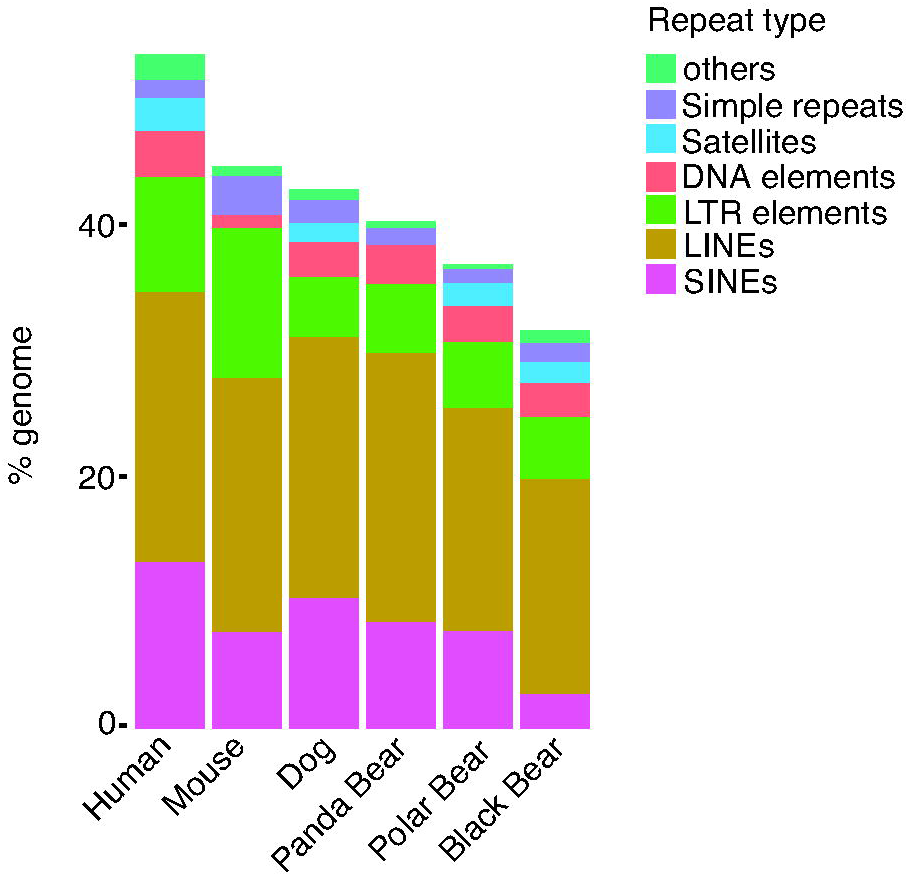
Comparison of different types of repeats between the black bear and several mammalian species shows a lower number of repeat sequences in the black bear.

### 3.2. Annotation of the Black Bear Genome

Our black bear genome assembly was annotated using the Comparative Annotation Toolkit (CAT). This toolkit uses whole genome alignments generated by progressiveCactus to project gene annotations from a high-quality reference genome on to one or more target genomes. This process leverages previously curated annotation sets to rapidly construct a set of orthologs in the target genomes. After transcript projection, the protein coding projections are provided to the *ab-initio* gene finding tool AUGUSTUS with additional extrinsic information derived from RNA-seq. AUGUSTUS then enforces a coding gene model that allows for shifts in splice sites in order to maintain frame. This process can also rescue exons that did not align in the whole genome alignment.

CAT identified 29,624 genes (18,091 coding) in our black bear assembly, representing 88% of the genes present in the dog annotation. In addition, 2,995 transcripts with at least 2 splices were predicted by AugustusCGP, which also had at least one intron junction supported by RNA-seq and not supported by transMap. Of these, 1,730 were associated with a transMapped gene and assigned as a novel isoform of that gene. The remainder were evaluated for being putatively novel loci. 653 loci were identified as being possibly paralogous, which is defined as overlapping a paralogous transMap projection, which was dropped in paralog resolution. 28 loci were identified as possible false fusions, defined as having >80% overlap with more than one transMap projection. 458 loci were identified as poor alignments, which are predictions in regions, which have alignment to a known gene but did not pass transMap filtering. This left 126 putatively novel loci. Of the 22,081 orthologous protein coding transcripts identified, pairwise codon-aware alignments of coding sequences identified frame-shifting indels in 7,848 transcripts. A slight enrichment of frame-shifting deletions was seen relative to insertions, suggesting a systematic bias in the assembly process.

### 3.3. High-throughput Sequencing of Renal RNA and Differential Expression Between Fall and Spring

Kidney samples were collected in the fall, before hibernation, and in the spring, shortly after the bears emerged from hibernation (3 males and 3 females for each time point). Approximately 60 million RNA-Seq reads were obtained from each sample before quality control (Table S5). After appropriate quality control and correction for batch effect, we performed a principal component analysis (Supplemental Fig. 1a) and determined that sample 101 (from a male in the spring) was an outlier. After removing this sample and repeating the analysis (Supplemental Fig. 1b) we compared the spring samples to the fall samples. We identified 169 differentially expressed protein-coding genes with an adjusted P-value below 0.05 (Fig. 3 and Table S6). Of these, 101 genes were downregulated and 68 genes were upregulated after hibernation. Fold changes ranged only from 1.8-fold downregulation (*RTN4RL2*) to 2.4-fold upregulation (*CISH*). Figure 4 shows a heatmap for all 11 bears with the 50 most significant genes (Supplemental Fig. 2 shows all 169 genes). Pathway enrichment analysis (Ingenuity Pathway Analysis) of the 169 genes did not identify any specific pathways. However, most notable was the upregulation of three cytokine suppression genes (*SOCS2, CISH,* and *SERPINC1*) and the lack of increased cytokine expression (e.g. *IL6, CCL2, CCL6*) and damage markers (*LCN2* and *HAVCR1*) normally seen in lower functioning or recovering kidneys of other species ^26^.

**Fig. 3.**
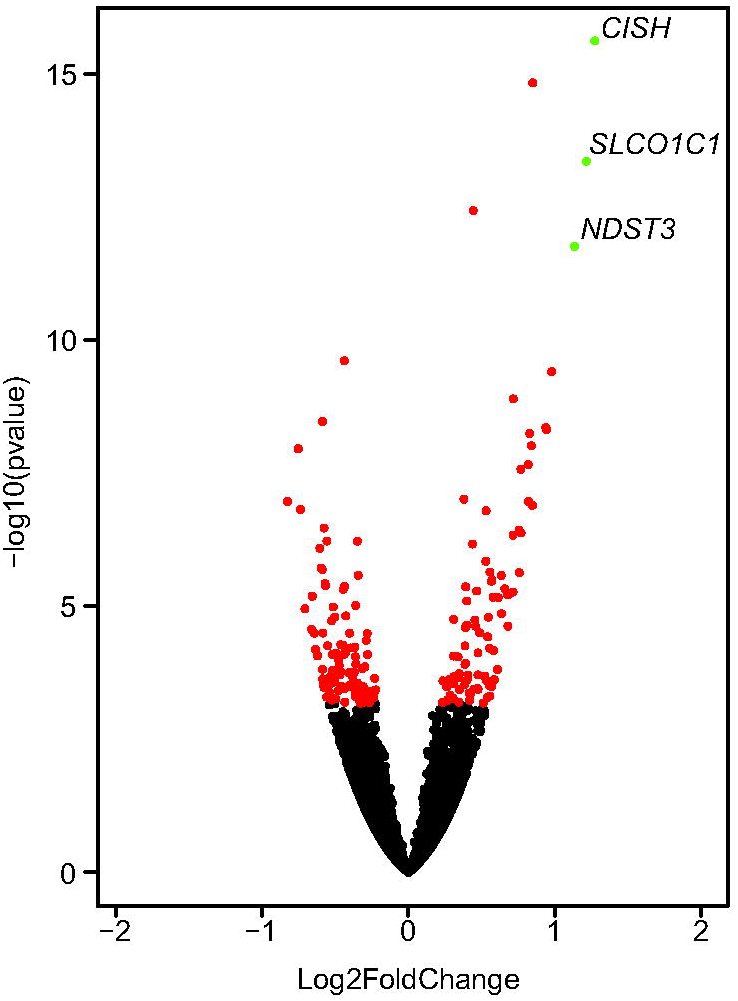
Volcano plot of the differential gene expression between spring and fall with *CISH, SLC01C1,* and *NDST3* being upregulated > 2-fold in the spring compared to the fall. All genes (169) with an adjusted P-value below 0.05 are indicated in red.

**Fig. 4.**
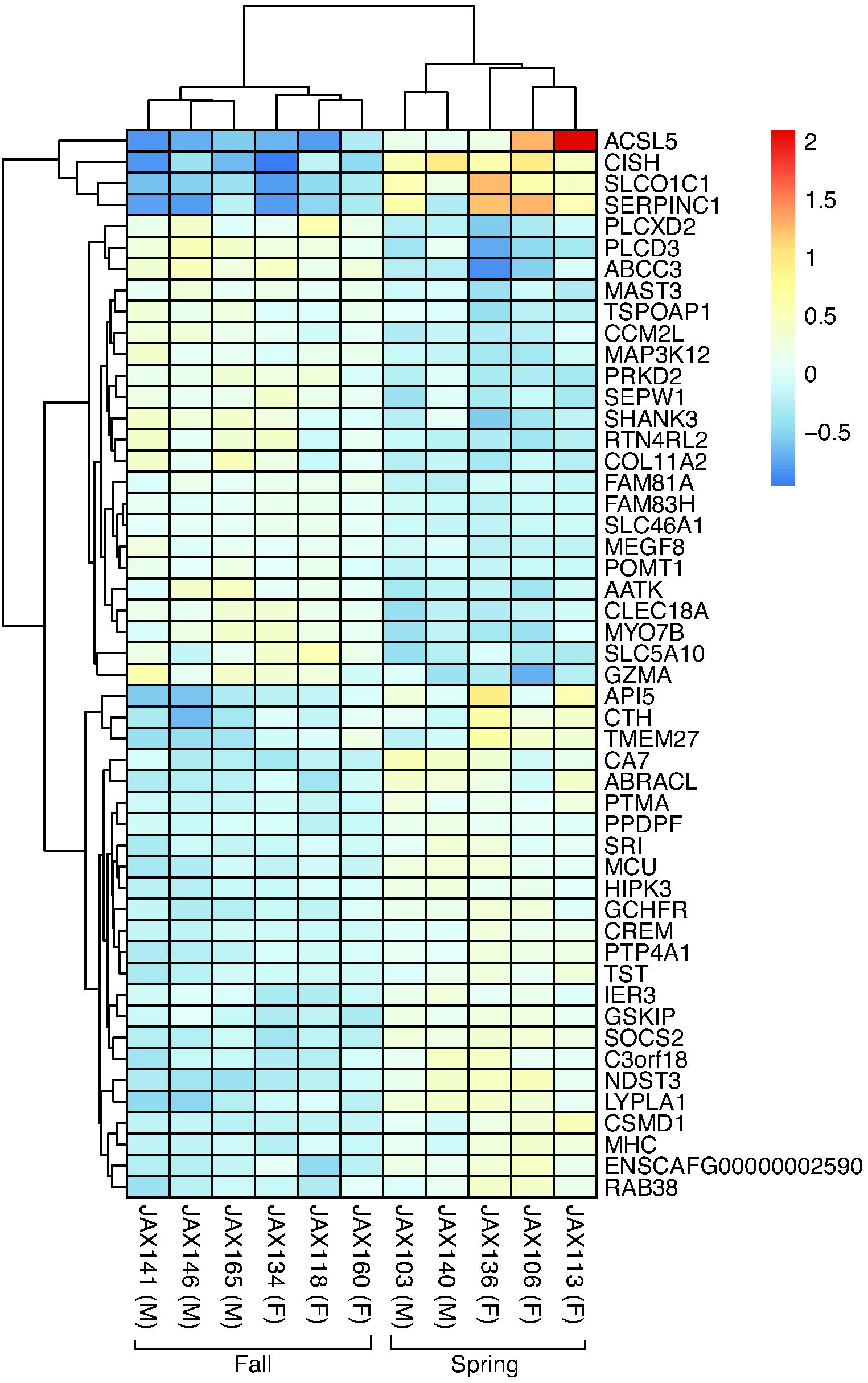
Cluster analysis of the gene expression profiles showing the 50 genes with the lowest P-value.

Because of the lack in inflammatory response, we asked whether the differentially expressed genes matched a particular development stage of the kidney. We used a method previously employed to place human cancers on a developmental landscape ^27^. First, we obtained gene expression profiles from mouse kidneys at different developmental stages (from E12.5 through E16.5) that were deposited in NCBI GEO and plotted the developmental expression profile in their first two temporal principal components (PC1-2). The high-dimensional expression profiles are simplified into a developmental timeline, ordering the genes in a linear array based on their temporal pattern of expression. Early genes are localized on the left end, genes with no bias towards early or late expression center in the middle and late genes localize the right end. Thus, the unique order of genes on the timeline represents a summary of early and late states for each developmental process. Next, we matched the differentially expressed bear genes with their mouse orthologs (153 out of 169 genes) and plotted their location on the developmental timeline for the genes that were expressed in the developing kidney (109 genes). We observed clustering of downregulated genes with the early mouse genes (E12.5) and clustering of upregulated genes with the late mouse genes (16.5) (Figure 5). The processes that take place in the E12.5 kidney are focused on progenitor cell dynamics, FGF, and Wnt signaling, which we know are pro-fibrotic in the adult. In contrast, E16.5 is very different as it involves glomerulogenesis, tubulogenesis and maturation.

**Fig. 5.**
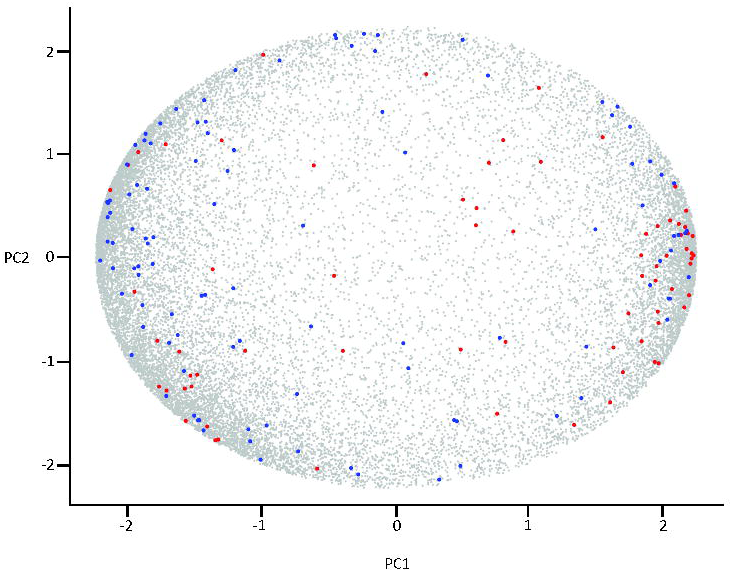
The mouse kidney developmental expression profile at E12.5-E16.5 in their first two temporal principal components (PC1-2) with the mouse orthologs of genes that are downregulated after hibernation in blue and genes that are upregulated in red.

Taken together, despite the many months of reduced renal function and down-regulated metabolism during hibernation, the gene expression differences between spring and fall bears is limited both in number of genes that are differentially expressed, fold-change, and the lack of upregulation of genes that are typically seen in other species after a period of low function or damage. In fact, the expression profile of bears coming out of hibernation resembles a developmental stage.

### 3.4. Identification of RNA-editing in the Black Bear Kidney

Our RNA-Seq data identified possible RNA-editing in a small set of genes when aligning the reads to our assembled genome sequence. In order to rule out the possibility of variation in the transcripts due to single nucleotide polymorphisms (SNPs) in the genomes of the bears for which we performed the RNA-Seq, we sequenced the genomes of bears 101, 103, 113, 118, 134, and 160 at a low coverage (Supplemental Table 7). After comparing with the genome sequences and filtering for protein-coding transcripts we identified 38 transcripts containing a different variant (between 10% and 95% of the total reads) from the genome sequence (Supplemental Table 8). Of these 38 transcripts, 30 (79%) show A-to-I editing, while the other sites have C-to-U editing. Almost half of the edited sites are in the coding region and five are predicted to lead to an amino acid change.

According to the REDIportal (http://srv00.recas.ba.infn.it/atlas/) RNA-editing has been observed in four of the genes in the human kidney (although not at the same base pair) and for one gene, *FLNB*, RNA-editing also occurs both in the human and the mouse kidney. Interestingly, RNA-editing in 37 of the transcripts is observed in most or all 12 bears, but for *ZNF688* we find editing in all 6 spring bears and no editing in the 6 fall bears.

To confirm our observations in the high-throughput sequencing data, we randomly selected eight editing sites and designed primers flanking these sites that could discriminate between RNA and possible genomic DNA contamination. Subsequent Sanger sequencing confirmed all eight editing sites (Supplemental Table 8).

## 4. Discussion

Previous studies in bears suggest they have unique features in the kidney that allow them to endure lower renal functioning during hibernation and recovery soon after hibernation. As these features are likely in part encoded in the genome sequence and gene expression patterns unique to the bear, we set out to explore the genome sequence and renal gene expression of the American black bear (*Ursus americanus*).

A first important step was to establish the complete annotated genome sequence. In order to accomplish this, we used three approaches for sequencing a single male bear with 100x coverage before filtering. This resulted in a final, high quality, assembly. This current genome is comparable in quality and size to the published panda genome^24^ and polar bear genome.^23^ A comparison between the three genomes shows a narrower range in GC-content in the black bear (Fig. 1c), likely the result of fewer repetitive sequences caused by transposon integration (Fig. 2).

The new Comparative Annotation Toolkit (CAT) pipeline allowed us to establish a high-quality annotation of the genome with 22,081 protein coding transcripts and identification of 126 putative novel loci that warrant further investigation as to their uniqueness and possible function.

Based on the reported changes in the bear kidney function during hibernation,^5^ we expected we would find a signature of these changes in the gene expression of kidneys collected soon after hibernation when compared to kidneys collected before hibernation. We therefore collected samples from 3 male and 3 female bears (samples were collected within 10 days from each other) soon after hibernation and 3 male and 3 female bears before hibernation. One limitation is that the time between waking up from hibernation and sample collection is unknown and we assumed it was similar for all animals. Deviation from this assumption is expected to lead to more variation in the gene expression profile. RNA-Seq analysis identified 169 protein coding genes that were differentially expressed between the two time points. Of these, 101 genes were downregulated and 68 genes were upregulated after hibernation. Fold changes only ranged from 1.8-fold downregulation (*RTN4RL2*) to 2.4-fold upregulation (*CISH*) and only three genes showed a >2-fold change (*CISH, SLCO1C1*, and *NDST3*). Little is known about the function in the kidney of these genes: CISH is a member of the SOCS family and involved in suppression of cytokine signaling through the JAK-STAT5 pathway. SLCO1C1 is a member of the organic anion transport family and best known for mediating sodium-independent uptake of thyroid hormones in brain tissue. NDST3 is a member of the heparan sulphate / heparin GlcNAc N – deacetylase / N-sulfotransferase family and increased expression indicates an increase of N-sulfation of heparin sulfate in the kidney. Since kidney glucosaminoglycans have been shown to be related to the ability of turtles to conserve water,^28^ it can be speculated that the increased expression of NDST3 in hibernating black bear kidneys protect the animals during the long period of dehydration during winter sleep. Moreover, since the endothelial glycocalyx on the vascular endothelial cells plays a major role in albuminuria and kidney disease,^29^ it could be speculated that upregulation of NDST3 may protect the endothelial glycocalyx in bear kidneys during the anuric hibernation.

Most notable among the differentially expressed genes was the upregulation of cytokine suppression genes (*SOCS2, CISH,* and *SERPINC1*) in the post-hibernating bears and the lack of increased cytokine expression (such as *IL6* and *CCL2*) and damage markers such as *HAVCR1* (KIM-1) and *LCAN* (NGAL). Pathway enrichment analysis (IPA) did not identify any pathways that were significantly overrepresented among the differentially expressed genes. These findings support previous observation that despite several pro-inflammatory risk factors during the hibernation period (such as kidney dysfunction and immobilization), there are no signs of systemic inflammation ^30^. This is further supported by our principal component analysis in which the genes that are downregulated after hibernation cluster with a developmental stage in which FGF and Wnt signaling, which we know are pro-fibrotic in the adult kidney, play an important role and genes that are upregulated after hibernation cluster with a developmental stage in which glomerulogenesis, tubulogenesis and maturation are important. In patients with CKD, systemic inflammation is a common finding that promotes premature aging and predicts poor outcome ^31^.

Although this is the first study in which gene expression is analyzed in kidney tissue, it is not the first in which gene expression in bears has been examined. Fedorov *et al.* analyzed samples from captive bears that were either kept in hibernating or non-hibernating conditions with a custom microarray. They analyzed heart and liver samples and found 245 differentially expressed genes in the heart and 319 genes in the liver ^32^. We were able to map the majority (85.2%) of their ESTs onto our genome assembly and match them to our transcripts. However, none of the genes that were found to be differentially expressed in heart and liver overlapped with differentially expressed genes in the kidney. The lack of overlap could be due to difference in experimental conditions such as the timing of tissue harvest. Alternatively, it may indicate that while nearly every tissue adapts to hibernation, the ensemble of genes involved is highly tissue-specific.

Overall, both the number of differentially expressed genes and the fold-changes are low, but are in line with the findings of Federov *et al*. Although it has been shown that American black bears maintain a reduced metabolic rate for up to three weeks after emerging from their dens ^3^, a possible explanation for our results is that the kidneys already completely recovered from hibernation within the short time between emerging from hibernation and sample collection. Another explanation is that the reduction in metabolic rate and urine production has a much smaller impact on the kidney than would be predicted.

Our RNA-Seq data also identified the presence of RNA editing, which was confirmed for all sites that were tested with Sanger sequencing. Several of the genes that are edited also show RNA-editing in mouse and human, but at different positions (Supplemental Table 8). The most interesting is the RNA editing in the 3’UTR of the transcriptional regulator *ZNF688*, which only seems to happen in the spring. This suggests season-dependent RNA-editing and a possible mechanism through which gene expression is altered.

In conclusion, we established a high-quality and well-annotated genome sequence of the black bear (*Ursus americanus*). RNA-Seq of kidney samples before and after hibernation shows RNA-editing and the modest differential expression of a set of 169 genes that might be involved in the unique stress response due to hibernation in the bear. Our results suggest that during hibernation, changes in gene expression favors anti-inflammatory pathways. Further studies are needed to understand the effects of these expression differences in the kidney.

## Availability of data

All raw and analyzed data is publicly available as described in the Materials and methods sections above.

## Acknowledgements

The authors thank Heidi Munger, Ryan Lynch, Edith Schriever, Lucy Rowe, and Xiaoan Ruan, for technical assistance, and Carol Bult, Peter Stenvinkel, Leif Oxburgh, and Paul Shiels for discussion and critical review of the manuscript. A special thanks to the Penobscot Indian Nation and Passamaquoddy Tribe for providing bear kidney samples.

